# DNA metabarcoding-based detection of non-indigenous invertebrates in recreational marinas: influence of sample type and seasonal variation

**DOI:** 10.1101/2024.01.25.577180

**Authors:** Ana S. Lavrador, Fábio G. Amaral, Jorge Moutinho, Pedro E. Vieira, Filipe O. Costa, Sofia Duarte

**Affiliations:** Centre of Molecular and Environmental Biology (CBMA) and ARNET-Aquatic Research Network, Department of Biology, University of Minho, Campus de Gualtar, 4710-057 Braga, Portugal; Institute of Science and Innovation for Bio-Sustainability (IB-S), University of Minho, Campus de Gualtar, 4710-057, Braga, Portugal

**Keywords:** DNA metabarcoding, non-indigenous species, eDNA, settlement plates, zooplankton

## Abstract

Monitoring of marine invertebrate non-indigenous species (NIS) using DNA metabarcoding can be strongly affected by selected sample type due to life history traits, such as habitat preferences and life cycles. Two marinas in the north of Portugal were sampled to assess the impact of sample type (hard and artificial substrates, water eDNA, and zooplankton) and season (spring, autumn, winter) on species and NIS recovery. Using two molecular markers - the mitochondrial cytochrome c oxidase subunit I (COI) and the small subunit ribosomal RNA (18S) - a total of 636 species and 31 NIS were detected. Species numbers were slightly higher in the marina more exposed to maritime traffic, and the highest percentage of exclusive species was detected in zooplankton (up to 24%), as well as the highest numbers of NIS. Regarding season, the highest numbers of species and NIS were detected in the spring and autumn (varying within each marina). Taxonomic composition analysis revealed differences in species richness and community structure among seasons and sample types, particularly between hard and artificial substrates *versus* eDNA and zooplankton. Of the 31 NIS detected, six are potential first records for Portugal, which await morphology-based validation. No NIS were detected in all sample types nor in all sampled seasons. This highlights the need to employ different sampling approaches and markers, as well as consider seasonal variation and level of exposure to maritime-driven introductions to guarantee a comprehensive metabarcoding-based surveillance of NIS in recreational marinas.

## Introduction

Coastal ecosystems are regions of great ecological and socio-economic importance and a source of biological productivity, as well as of goods and services (Costanza et al. 1997). However, these are vulnerable regions, heavily impacted by human activities that often lead to loss of biodiversity (Turner and Rabalais 1994; Jackson et al. 2001). These ecosystems are also particularly susceptible to the introduction and spread of non-indigenous species (NIS).

Non-indigenous species are species that are introduced (naturally or by human mediation) outside their natural occurrence range. If these species successfully establish, reproduce, and spread, they can become invasive, posing a threat to the ecosystem’s integrity (Bax et al. 2003; Katsanevakis et al. 2014; Rilov and Crooks 2009). The introduction of NIS has become so frequent and relevant with the increase of globalization, that they are now included in important legislations and directives such as the European Union Regulation 1143/2014 on Invasive Alien Species and the Marine Strategy Framework Directive (European Comission 2008; European Union 2014; Diagne et al. 2021). It is also a concern for the conservation of oceans and marine resources (addressed in the UN Sustainable Development Goal SDG14), as invasive marine species can cause severe disruptions in native biodiversity, alter habitats, and compromise the delicate balance of marine environments, threatening the health and resilience of marine ecosystems.

One of the most significant vectors of NIS introductions in coastal ecosystems is shipping. Shipping vessels can transport NIS through biofouling (species that attach themselves directly to the ship’s surfaces, as well as species that live amongst those communities), and in the ballast waters (that are loaded into the ship to adjust buoyancy and secure stability and manoeuvrability, carrying an array of different species and life-forms) (Rilov and Crooks 2009; Sardain et al. 2019). Recreational marinas and harbours are, thus, important points of entrance for marine NIS, as they are recipients of different types of shipping vessels, harbouring several physical structures (e.g., pontoons, cables, buoys, anchors) that allow colonization and spread of fouling organisms (Glasby et al. 2007). With the increment in globalization and transport of goods and services by shipping (which is estimated to continue to increase up to 240-1209% by 2050) it is important to prioritize NIS surveillance in harbours, recreational marinas, and vicinity areas (Sardain et al. 2019; Bailey et al. 2020).

Most NIS-monitoring programs rely on the observation and morphological identification of species (Chainho et al. 2015; Mancinelli et al. 2017; Afonso et al. 2020). However, with the decline in the number of taxonomists worldwide, and challenging species identifications in aquatic systems (e.g., due to low visibility, cryptic taxa, low-density populations and life stages not amenable to morphological identification) (Knowlton 1993; Hopkins and Freckleton 2002; Kim and Byrne 2006), DNA-based methods emerged as complementary or alternative approaches. These techniques allow for the simultaneous processing of a large number of samples, and accurate identification of multiple taxa from diverse environmental samples, with increased sensitivity and specificity. They often reveal hidden diversity, while allowing greater time and cost effectiveness (Holman et al., 2019; Pochon et al., 2015; Rey et al., 2020; Taberlet et al., 2012). In particular, DNA/eDNA metabarcoding, that combines amplicon barcoding with high-throughput sequencing (HTS), is a powerful tool for identifying multiple species from complex samples (bulk organismal samples, environmental samples) (Hajibabaei 2012; Taberlet et al. 2012; Cristescu 2014). With this approach, accurate identifications of NIS adult, larvae and propagules can be obtained using standardized DNA barcode markers that target a wide taxonomic range of organisms in mixed samples. To that end, a partial segment of the mitochondrial cytochrome c oxidase subunit 1 (COI) gene has been established as a standard marker for monitoring metazoan diversity, providing reliable identifications, as well as the discrimination of closely related species (Hebert et al. 2003). Concurrently the small subunit ribosomal RNA gene (18S) has been widely employed in marine biomonitoring studies to detect eukaryotes, including macroinvertebrates (Pawlowski et al. 2018). DNA metabarcoding also allows the early detection of invasive species before its presence and impact become irreversible, and the increase of sampling density, compared to morphotaxonomic assessments, ultimately improving NIS monitoring in coastal ecosystems (Pochon et al. 2015; Holman et al. 2019).

The majority of European marine NIS are invertebrates (64.2% in 2020) (Zenetos et al. 2022). Marine invertebrates are a very diverse group of organisms, composed of many different phyla with distinct life cycles (i.e., with or without larval stage) and habitat preferences (i.e., benthic, pelagic, fouling organisms). As such, monitoring marine invertebrates requires various types of samples, as well as temporal and spatial assessments to recover a wider range of species and more accurately describe the communities present in coastal environments (Lehtiniemi et al. 2015; Koziol et al. 2018; Kraus et al. 2019). This is also important when monitoring non-indigenous invertebrate species as organisms introduced by hull fouling can pass to the hard artificial substrates of the marinas and harbours, through hull cleaning operations; or from the water column through larval stages or propagules (European Environment Agency 2021). Recent studies have shown that recreational marinas are more relevant for the spreading of NIS than it was supposed, as they present communities that diverge from those recovered in the nearest harbours, and some even present a higher number of NIS, or different NIS than those detected in the harbours (Ferrario et al. 2017; Chebaane et al. 2019; Afonso et al. 2020; Png-Gonzalez et al. 2021). This suggests that recreational vessels, and not only larger industrial vessels, are very relevant vectors for NIS spreading (Ferrario et al. 2017; Chebaane et al. 2019; Afonso et al. 2020; Png-Gonzalez et al. 2021). This can also be explained due to the larger docking times of these vessels in the recreational marinas, as NIS spreading can vary with the origin of the vessels, as well as the duration of time moored to the marina (Martínez-laiz et al. 2019; Tempesti et al. 2022). Given the importance of the early detection and monitoring of marine invertebrate NIS in recreational marinas, and the role of season and sample type in NIS recovery through DNA metabarcoding, our specific aims were to test the effects of these factors using as a case study two recreational marinas in the north of Portugal. These marinas differ in maritime traffic: Viana do Castelo, located in a region of low to moderate maritime traffic, and Porto Atlântico, near the second highest maritime traffic harbour in Portugal. For this study, we analysed communities recovered from five different sample types: marinas’ hard substrates (i.e., pontoons, cables, buoys), artificial substrates (tri-dimensional sponges and acrylic plates), water for environmental DNA (eDNA), and zooplankton, in 3 seasons during 1 year (March 2020-March 2021), to determine its impact on macroinvertebrate species recovery, particularly NIS detection, through DNA metabarcoding.

## Results

### Initial HTS datasets

Two molecular markers were used for species identification through HTS. The total number of raw reads resulting from 90 datasets (three sampling points x two marinas x five substrates x three seasons) was 3.171,606 and 3.584,901 for COI and 18S, respectively. After primers sequences removal, and merging of forward and reverse reads in mothur, 65 and 75% of the reads were submitted respectively for analysis in mBRAVE and SILVAngs (2.067,067 reads for COI and 2.693,307 for 18S). After subsequent filtration steps, 2.025,812 (COI) and 2.570,633 (18S) usable reads remained for taxonomic classification, of which 811,044 (40%) and 486,252 (19%) were assigned to species, for COI and 18S, respectively. From these, 749,290 and 444,877 reads belonged to marine/oligohaline invertebrates, of which 15% and 10% were assigned to non-indigenous species (NIS) (for COI and 18S, respectively) (Table S1). Overall, 18S recovered more species than COI. Given that a high percentage of species was detected exclusively by each marker, the remaining analyses were performed using the combined information from both markers. In this study, a total of 636 invertebrate species were recorded and 31 NIS detected.

### Effect of sample type in total species and NIS diversity

Five types of samples were assessed to study its effect on the detection of total species and NIS in the two recreational marinas: organisms fouling hard substrates of the marina, organisms colonizing sponges and acrylic plates (artificial substrates), eDNA from the water, and zooplankton. In both marinas, considering the combined data from the three seasons, zooplankton was the strategy that recovered the highest numbers and percentage of exclusive species (17 to 24%), followed by hard substrates and eDNA, with plates recovering the lowest numbers and percentage of exclusive species (up to three percent) (Fig. 1A and B). Individually, zooplankton recovered 67 to 59% of all species detected; sponges 47 to 33%; eDNA 46 to 40%; hard substrates 44 to 41%; and plates 33 to 27%. eDNA and zooplankton were the sample types sharing the highest percentage of species (six to eight percent), followed by zooplankton and hard substrates in Viana do Castelo (three percent) and hard substrates and sponges in Porto Atlântico (four percent), while the percentage of shared species was generally low among other pairs of sample types (zero to two percent).

**Fig. 1.**
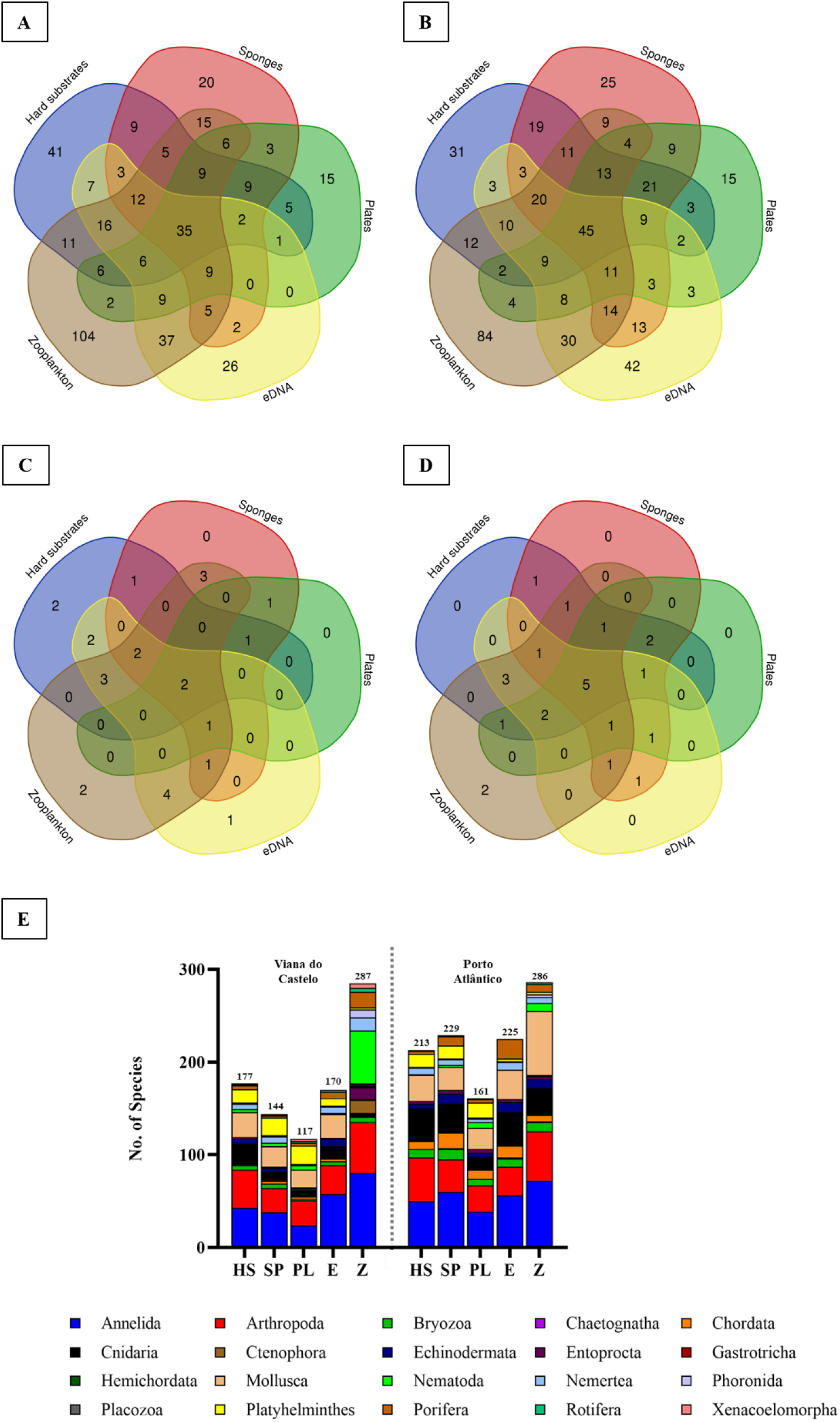
Total number of species (**A** and **B**) and NIS (**C** and **D**) detected in the three sampling seasons for each sample type, in the Viana do Castelo (**A** and **C**) and Porto Atlântico (**B** and **D**) recreational marinas. **E** Taxonomic composition of the total marine invertebrate species detected in the entirety of this study in the Viana do Castelo and Porto Atlântico recreational marinas, discriminated by sample type. HS: hard substrates. SP: sponges (artificial substrate). PL: acrylic plates (artificial substrate). E: eDNA. Z: zooplankton.

Regarding NIS, zooplankton recovered the highest numbers in both marinas (along with hard substrates in Porto Atlântico), and the plates recovered the lowest number of NIS (Fig. 1C and D) (Table S2 and S3). When considering the overlap of species and NIS detected by all sample types, only eight and nine percent of the total species and eight and 20.8% of NIS (for Viana do Castelo and Porto Atlântico, respectively) were detected in common using all the sample types (Fig. 1).

Regarding the taxonomic composition of these communities, in general most species detected in both recreational marinas belonged to Annelida and Arthropoda (Fig. 1E). Among sample types there were some notable differences particularly when comparing hard and artificial substrates *versus* eDNA and zooplankton samples. Overall, in hard and artificial substrates there were more species belonging to Platyhelminthes; less Porifera; less Nemertea in Viana do Castelo; and less Annelida and Arthropoda, when compared to eDNA and zooplankton. In contrast, there were more Nematoda, Phoronida and Rotifera species in eDNA and zooplankton samples and Mollusca species in zooplankton were detected in fewer numbers in Viana do Castelo, while in Porto Atlântico they were highly present. Overall, there were very few Chaetognatha, Gastrotricha and Hemichordata species identified in the entirety of the study, but these few records were mostly recovered in zooplankton samples. Mollusca species were always detected in higher numbers in eDNA samples and zooplankton in Porto Atlântico (Fig. 1E).

When analysing the structure of communities among sample types, there is a clear separation between communities recovered in the hard substrates, sponges and plates *versus* communities recovered in eDNA and zooplankton, on each recreational marina (with the exception of eDNA samples from T2 in Viana do Castelo) (Fig. 2). The PCoAs also show distinct communities recovered in the winter, compared to summer and autumn samples in Viana do Castelo. The PERMANOVA analysis supported these conclusions, as the species composition recovered on each recreational marina was significantly influenced by sample type and season, and the interaction between these factors (P=0.001, for all factors, in both marinas) (Table S4).

**Fig. 2.**
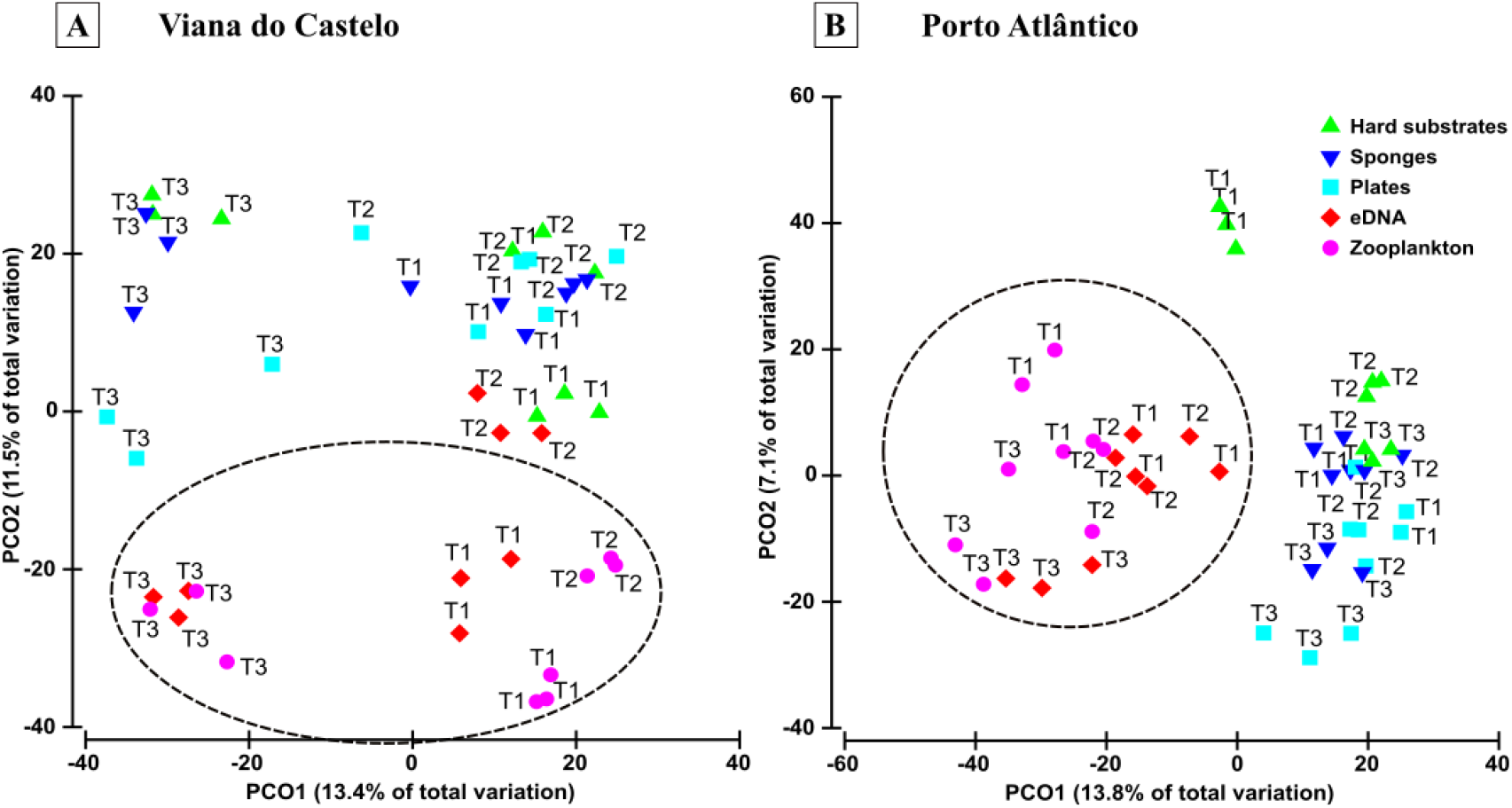
Principal coordinate analysis (PCoA) of sampled communities in Viana do Castelo (**A**) and Porto Atlântico (**B**) recreational marinas, in order to analyse the effects of season and sample type on community structure. Hard substrates: organisms fouling the hard substrates of the marina. Sponges: organisms colonizing 3-dimensional sponges deployed in the marinas. Plates: organisms colonizing acrylic plates deployed in the marinas. eDNA: environmental DNA extracted from water. Zooplankton: zooplankton collected from the water column. Black dotted line circles group eDNA and zooplankton samples.

### Effect of season in total species and NIS diversity

Regarding season, and combining data from all sample types, the highest numbers of species were detected in the spring (284) and autumn (348) (for Viana do Castelo and Porto Atlântico, respectively) and the lowest was detected in the winter, in both marinas. Only 11.3 to 24% of species were detected exclusively on each season. In terms of NIS, the highest numbers were detected in the autumn (21), for Viana do Castelo, and in spring (21), for Porto Atlântico, while the lowest were observed in the winter in Viana do Castelo and in the autumn and winter in Porto Atlântico (Fig. 3). Spring (Viana do Castelo) and autumn (Porto Atlântico) were the seasons that recorded the highest numbers of exclusive species (24 and 19.5%, respectively), while autumn was the season detecting the highest number of exclusive NIS in Viana do Castelo (23%) while in Porto Atlântico all seasons detected one NIS exclusively (4%). Spring and autumn were the seasons sharing the highest percentage of invertebrate species in both marinas (14 to 20%). In what respects NIS, spring and autumn were the seasons sharing the highest percentage for Viana do Castelo (27%), while the highest percentage of species shared for Porto Atlântico was found between spring and autumn as well as spring and winter (12.5%).

**Fig. 3.**
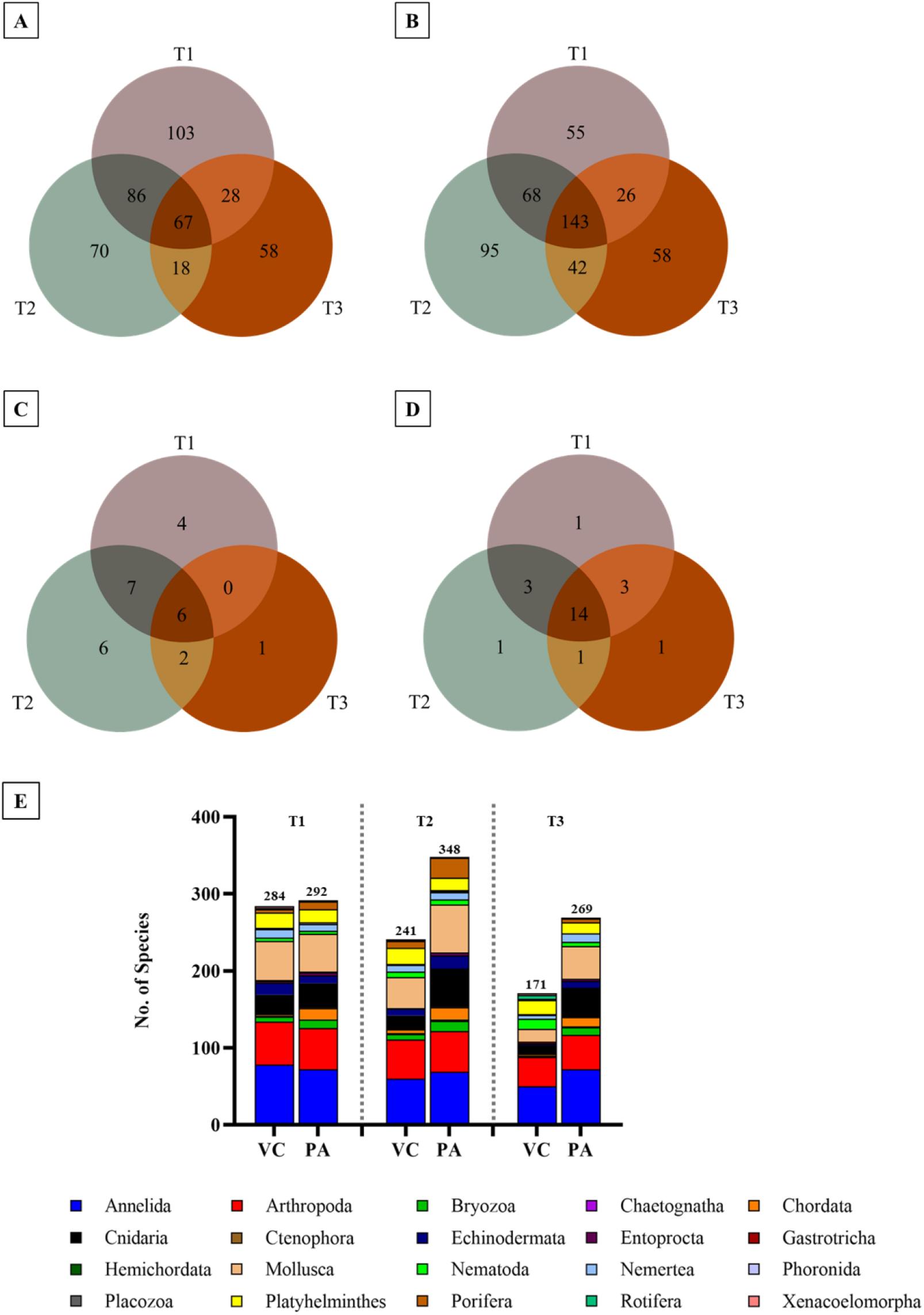
Total (**A** and **B**) and non-indigenous species (**C** and **D**) detected during the three sampling seasons, in the recreational marinas of Viana do Castelo (**A** and **C**) and Porto Atlântico (**B** and **D**). **E** Taxonomic composition of the total marine invertebrate species detected by season in the recreational marinas of Viana do Castelo (VC) and Porto Atlântico (PA).

Regarding the variation of the taxonomic composition of the communities within each season, in Viana do Castelo, in the spring and autumn, a very similar species composition was detected, with a dominance of Arthropoda and Annelida, however Mollusca decreased sharply in winter. The number of species belonging to Nematoda increased from spring to winter, while Echinodermata had a pronounced decrease after spring. In the winter, most species belonged to Annelida, Arthropoda and Platyhelminthes; however, there was an increase in the number of species belonging to Rotifera, in comparison with the other seasons. In Porto Atlântico, Arthropoda, Annelida, and Mollusca were the dominant groups in all seasons. From spring to autumn, there was an increase in the number of species of Cnidaria and Porifera, that then decreased from autumn to winter (Fig. 3E).

### Effect of the location in total species and NIS diversity

In Viana do Castelo, in total, 430 species and 26 NIS were recovered, while in Porto Atlântico, 487 species and 24 NIS were identified (Fig. 4; Table S2 and S3). Forty four percent of species and 61% of NIS were detected in common in both marinas, respectively.

**Fig. 4.**
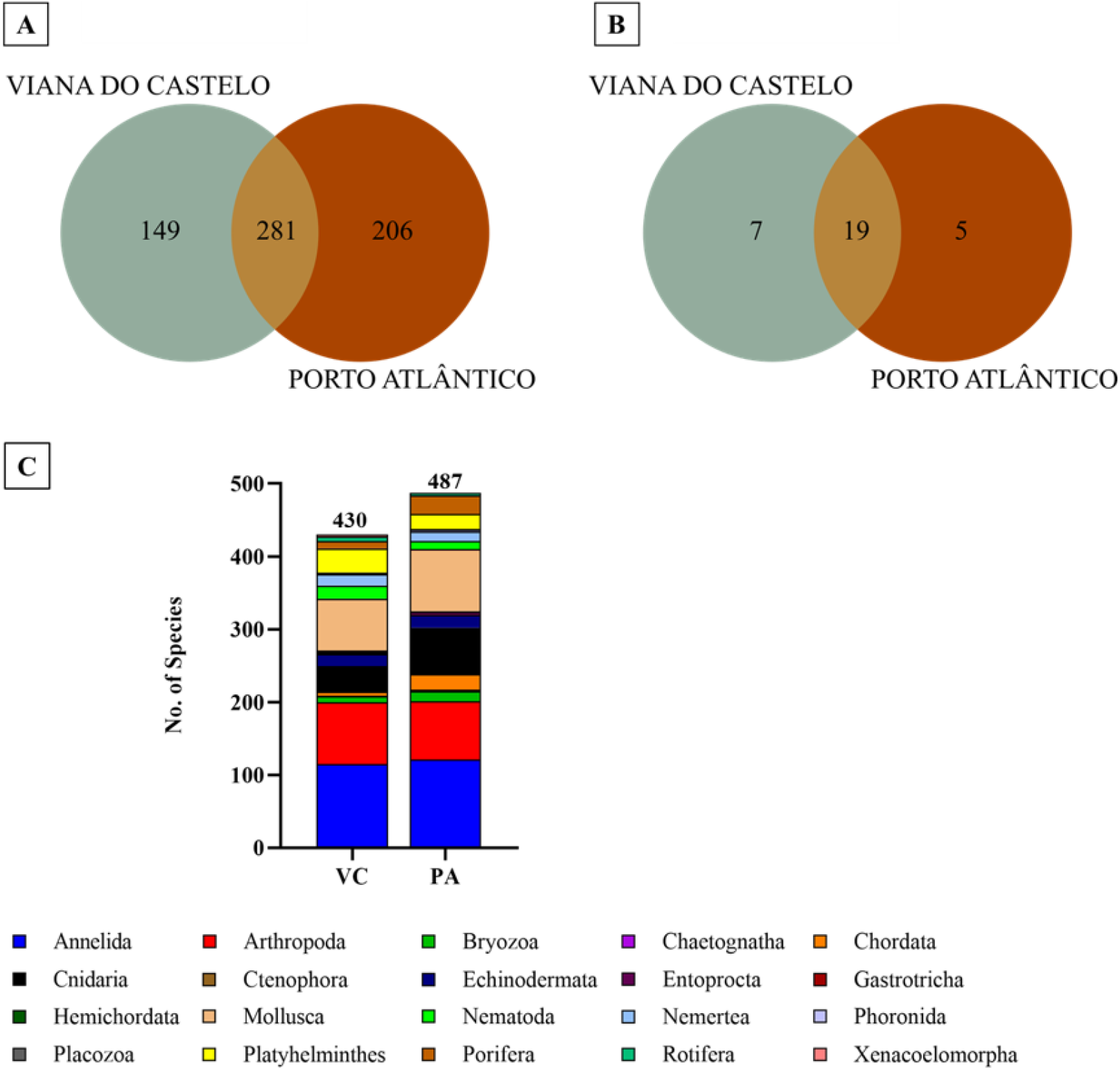
Total (**A**) and non-indigenous species (**B**) detected during the three sampling seasons and using all sample types, in the Viana do Castelo and Porto Atlântico recreational marinas. (**C**) Taxonomic composition of the total marine invertebrate species detected in the entirety of the study in the Viana do Castelo (VC) and Porto Atlântico (PA) recreational marinas.

Globally, the distribution of representatives of major invertebrate groups was similar among the two marinas (Fig. 4), despite some specific differences. When considering data from the three seasons combined, in both marinas, the majority of species belonged to Annelida, Arthropoda, and Mollusca. In Porto Atlântico we recorded a higher percentage of Chordata (Ascidiacea) and a lower percentage of Rotifera, when compared to Viana do Castelo. There were also more Bryozoa, Cnidaria and Porifera species; while higher numbers of Nematoda, Nemertea and Plathyelminthes and Xenacoelomorpha were detected in Viana do Castelo.

### Non-indigenous species (NIS) detection through DNA metabarcoding

In this study a total of 31 marine invertebrate non-indigenous species were detected. The updated list of European NIS retrieved from public databases contained 1,333 species of marine/oligohaline invertebrates, of which 60 were also present in our final list of species that were detected in this study. The native distribution of these 60 species (European NIS) was analysed, resulting in a list of 31 NIS detected in this study that are non-indigenous in Portugal, of which 6 appear to be first records (Table S3). The majority of NIS detected belonged to Arthropoda (Thecostraca), Chordata (Ascidiacea) and Annelida (Fig. 5).

**Fig. 5.**
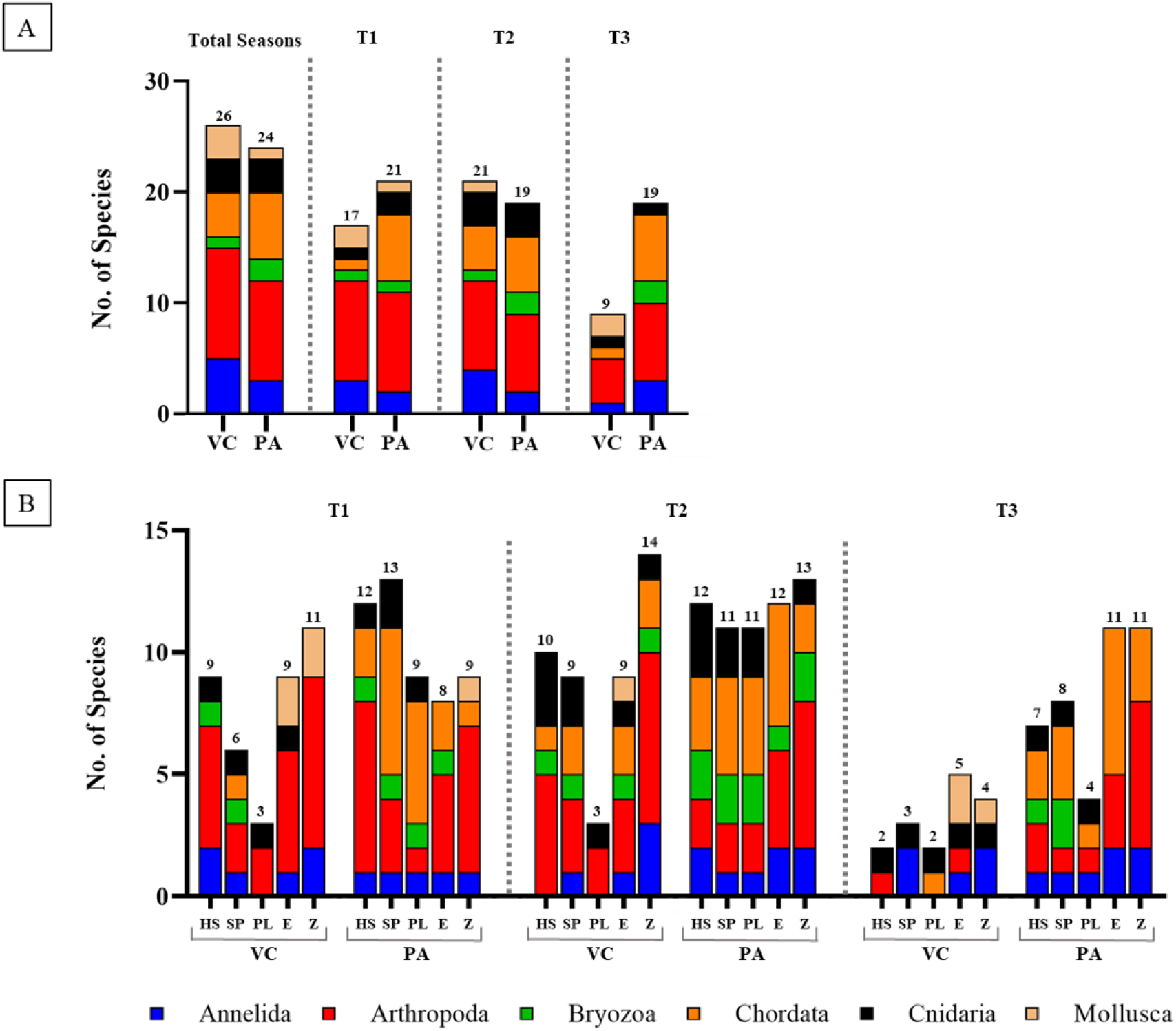
Taxonomic composition of the total non-indigenous marine invertebrate species detected in the entirety of this study in the Viana do Castelo (VC) and Porto Atlântico (PA) recreational marinas, discriminated by season (**A**) and per sample type, season, and recreational marina (**B**). T1: spring. T2: autumn. T3: winter. HS: hard substrates. SP: sponges (artificial substrate). PL: acrylic plates (artificial substrate). E: eDNA. Z: zooplankton.

These NIS were all identified by DNA metabarcoding (molecular identification). Given the importance of reliable identifications of species, in particular of non-indigenous species that can trigger governmental actions, all molecular identifications of NIS were further analysed. Of the 31 NIS, 12 were detected with COI exclusively, 11 with 18S exclusively, and 8 were detected by both markers (COI and 18S). 18S sequences for the 11 NIS exclusively detected with this marker were queried for a BLASTN search against the standard databases of NCBI, and only three molecular identifications were found to be reliable: *Eriocheir sinensis*, *Lovenella assimilis* and *Polydora onagawensis*. COI records were also further analysed, and of the 12 NIS exclusively detected with the COI marker, only five were records considered absolutely reliable: *Amphibalanus amphitrite*, *Austrominius modestus*, *Bugula neritina*, *Jassa slatteryi* and *Potamopyrgus antipodarum*. *Botrylloides violaceus* and *Lovenella assimilis* (detected exclusively with 18S) had also been identified using the COI marker. However, because the BINs that matched with these species were considered unreliable (e.g., also matched with other species), the species were removed from analysis with COI. Of these 31 NIS, 11 species are not suitable for morphological identification, as they were identified exclusively in eDNA or zooplankton samples; the remaining species were detected using sample types (hard and artificial substrates) that may allow further morphological verification, since bulk samples were again preserved in absolute ethanol, after DNA extraction.

Six NIS appear to be first records, at least to our knowledge, in Portugal: *Caprella mutica*, *Haliclystus tenuis*, *Lovenella assimilis*, *Oithona davisae*, *Phallusia nigra* and *Polydora onagawaensis*, recovered in different seasons and using different sample types. However, of these, only *Oithona davisae* had reliable matching records in the reference DNA databases (it was detected with both molecular markers and the BIN matching recovered COI sequences is concordant – see methods section for more details). *Musculus lateralis* was also detected in all three seasons and in both marinas (except for Viana do Castelo in the winter) and although its presence has not yet been confirmed in Europe and, thus, we did not consider it as a NIS, it has a great potential to be one since it has been documented only in North and Central America.

*Caprella mutica* was detected in Porto Atlântico in the spring and autumn (sponges) and winter (hard and artificial substrates). *Haliclystus tenuis* was detected only in the autumn in Viana do Castelo, but during the three seasons in Porto Atlântico (only in hard and artificial substrates). *Lovenella assimilis* was detected in the spring and autumn in hard substrates and sponges in Porto Atlântico; *Oithona davisae* in the first two seasons in Viana do Castelo (only in zooplankton, in spring, and in hard substrates, eDNA and zooplankton, in autumn); while in Porto Atlântico, it was detected in the spring (sponges and zooplankton) and winter (eDNA and zooplankton). *Phallusia nigra* was exclusively detected in Porto Atlântico, in sponges, in the spring and eDNA in the winter and, finally, *Polydora onagawaensis* was detected in Viana do Castelo in the first two seasons (hard substrates, eDNA and zooplankton, in spring, and, in zooplankton in autumn; and in Porto Atlântico in zooplankton in the winter. Overall, results found for each recreational marina show that no NIS were detected in all sample types and in all sampled seasons.

## Discussion

The major findings of this study can be summarized in four main points: 1) only eight to nine percent of species were detected in all types of samples, and only up to 24% of the species were detected exclusively in one sample type; 2) the lowest species richness was consistently recorded in the winter in both marinas, and the highest number of invertebrate species and NIS was either found in the spring or autumn; 3) more invertebrate species were recovered in Porto Atlântico (487 species and 24 NIS), a marina more exposed to maritime traffic and displaying higher salinity, but more NIS were recovered in Viana do Castelo (430 species and 26 NIS); and 4) a total of 31 marine invertebrate NIS were detected, of which 6 are potential first records for Portugal.

### Effect of sample type in total species and NIS diversity

One of the main goals of this study was to assess the influence of five different sample types – hard substrates, artificial substrates (sponges and plates), eDNA and zooplankton - in the recovery of marine invertebrate NIS in recreational marinas. To our knowledge, this is the first study using these five particular sampling strategies in the southwestern European coast employing molecular tools, and substantial differences were found in communities captured by hard and artificial substrates, particularly when compared to water eDNA and zooplankton. This is particularly relevant, since water eDNA has been the most employed for assessing NIS communities through DNA metabarcoding, in marine and coastal ecosystems (Borrell et al. 2017; Deiner et al. 2018; Grey et al. 2018; Rey et al. 2019; Wood et al. 2019; Suarez-Menendez et al. 2020; Duarte et al. 2021). In fact, and also supporting our conclusions, studies employing both water eDNA and bulk organismal samples suggest that eDNA alone cannot replace organismal sampling (e.g., Huhn et al., 2020; Leduc, Lacoursière-Roussel, et al., 2019; Rey et al., 2020; Westfall et al., 2020). Although dominant phyla found in our study were common to all substrates - Arthropoda, and Annelida (except for eDNA samples in Porto Atlântico, where Annelida and Cnidaria were dominant) - the percentage of species captured by all was significantly low – up to nine percent - and highlights the need to employ multiple sampling strategies to recover as much as possible the diversity in a given location through DNA metabarcoding.

This is in line with few previous studies employing multiple sampling strategies (e.g., Koziol et al., 2018; Rey et al., 2020) that concluded that different sample types recover different sets of species. This result is not totally unexpected, since marine invertebrates are highly diverse phylogenetically, consisting of species with very different traits such as mobility and attachment styles, feeding habits and colonization processes (Brusca et al. 2016). This makes it more difficult to study these organisms and their assemblages over time and space. Some organisms colonize surfaces forming biofouling communities that change over time, while others are benthic (living on the sea floor), or possess larval stages that inhabit temporarily the water column and may be mostly found in zooplankton, depending on the sampled season (Vinagre et al. 2020). In addition, and due to the presence of many artificial substrates (e.g., floating pontoons, cables, boats, buoys) and higher availability of nutrients, recreational marinas are highly prone to be colonized by biofouling communities (e.g., mussel lines) which *per se* can also constitute an augmented surface of potential niches available for NIS settlement (Png-Gonzalez et al. 2021).

In our study, plates recovered the lowest number of species and NIS – see Tables S2 and S3 - and communities in plates, as well as in sponges or hard substrates were dominated by Annelida and Arthropoda (regardless of season and marina). Previous studies aiming at assessing NIS diversity in hard substrates of Portuguese recreational marinas by using morphology-based approaches also showed a dominance of Arthropoda and Annelida (Afonso et al. 2020). In addition, a dominance of Arthropoda on PVC plates (deployed for approx. 3 months) was also previously found in the port of Bilbao (Spain) (Rey et al. 2020). But in a six-year survey using PVC plates deployed in a recreational marina in Madeira island (collected from 3 to 74 months), Bryozoa and Ascidiacea were the most represented groups (Canning-Clode et al. 2013). Also, Ascidiacea dominated plates deployed for 2 months in two coastal ports in Western Australia (Koziol et al., 2018).

On the other hand, zooplankton recovered the highest number of species and a few taxonomic groups were indeed predominantly found in this sample type, such as Nematoda, Phoronida and Rotifera, as well as a higher number of Annelida, Arthropoda and Mollusca species (in Porto Atlântico) and Nematoda (in Viana do Castelo). This is not totally unexpected since marine zooplankton community includes many different species of animals (i.e., from microscopic protozoans to large dimension animals), including holo- (spending their entire lives in the pelagic environment – e.g., Rotifera) and meroplanktonic species (i.e., eggs and larval stages of many benthic invertebrates’ species that are temporary members of the plankton).

Previous studies also employing DNA metabarcoding on multiple substrates reached conclusions similar to ours - each sampling method recovered a distinct set of organisms (Koziol et al. 2018; Pearman et al. 2021; Rey et al. 2020). And although comparisons among studies are hard to reach since sampling designs usually differ (i.e., sampling effort – including spatial and temporal) as well as the use of different methodologies along the DNA metabarcoding analytical chain (i.e., DNA extraction protocols, molecular markers and primers, bioinformatic pipelines), most studies have attained the same conclusion: (e)DNA metabarcoding is a very proficient tool for monitoring and identifying species and taxonomic profiling of communities. However, a sampling design combining multiple sample types, spatio-temporal variation, and the use of multiple molecular markers, may be needed for a more comprehensive surveillance of NIS in these systems. We support these conclusions as our study revealed that if only one sample type had been used exclusively to monitor species and NIS communities in these marinas, 33 to 73% of all species and 25 to 81% NIS would have gone unnoticed (depending on sample type and marina). Some of these examples include *Mercenaria mercenaria*, *Mya arenaria*, *Polydora cornuta*, *Potamopyrgus antipodarum* and *Streblospio benedicti* that were exclusively detected on zooplankton and/or eDNA samples, while *Blackfordia virginica*, *Caprella mutica*, *Haliclystus tenuis* and *Lovenella assimilis* were detected exclusively in hard and/or in artificial substrates.

### Effect of season in total species and NIS diversity

There were prominent differences in community composition of samples retrieved in the winter *versus* those retrieved in the spring and autumn in Viana do Castelo. This is expected as winter affects species richness and diversity due to its lower temperatures (Blomquist and Bonsdorff 1986). In marine invertebrates it has also a notable influence on its life cycle, for example, certain crustaceans and molluscs enter a state of dormancy or reduced activity due to the low environmental temperature, affecting their availability in the ecosystem (Boss 1974; Alekseev and Starobogatov 1996). In this period there is also a lower photoperiod which leads to limited food availability (Cloern and Jassby 2010) which combined with less shelter availability can force species to adapt or seek refuge in different habitats. The water temperature data collected during this study also supports these conclusions as it was very similar in all seasons, but significantly lower in the winter (Fig. S1). All these factors contribute to fluctuations in species composition and abundance. These differences due to seasonality further justify the need to sample broader time spans to better understand species, NIS communities and their dynamics and to further correlate NIS introductions with biotic and abiotic factors (Gittenberger et al. 2023).

### Effect of location on total species and NIS recovery

The two recreational marinas studied have particular geographic settings that can account for some of the differences observed. Viana do Castelo marina is located on the north bank of the Lima River estuary, it is more sheltered, and the water has a lower current flow (personal observation). Porto Atlântico marina is around 2.5 miles from the mouth of the Douro River and at the entrance of the Leixões trading seaport, which is the second largest in Portugal, in terms of maritime traffic. The trading ports closest to each of the recreational marinas also present different levels of vessels’ flow, which could justify the higher presence of species in Porto Atlântico marina. Data collected from each port from March 2020 to March 2021 shows that 746 vessels from 38 different countries were docked on the Leixões port, while Viana do Castelo harboured 203 vessels from 22 countries (approximately one quarter of Leixões vessels) (Fig. S2). Although NIS origins and introduction pathways were not in the scope of this study, considering that the main vector of NIS introductions in aquatic ecosystems in Europe is through shipping (both biofouling and ballast waters), it is important to analyse the marina’s vessels flow and permanence, as well as its origin (Katsanevakis et al. 2013). In our study, even though the Porto Atlântico marina was near a harbour with a higher flux of vessels that had a broader number of countries of origin, higher numbers of NIS were detected in Viana do Castelo. However, this difference in vessel flux can account for the higher number of invertebrate species found in Porto Atlântico. In the latter, communities seemed also to be more affected by the sample type than season, probably due to geographical differences of the marina, as well as of the different chemical parameters of the water (greater salinity in Porto Atlântico).

### Non-indigenous species (NIS) detection through DNA metabarcoding

In this study, a total of 31 NIS was detected by DNA metabarcoding, six of which are potential first records for Portugal. Eight of these NIS are also present in the National List of Invasive Species (by the Portuguese Institute for Nature Conservation and Forests): *Amphibalanus amphitrite*, *Austrominius modestus*, *Blackfordia virginica*, *Botryllus schlosseri*, *Corella eumyota*, *Mya arenaria*, *and Tricellaria inopinata*, as well as *Eriocheir sinensis* which is also listed as one of the 100 of the world’s worst invasive alien species by the International Union for Conservation of Nature (IUCN) (Lowe et al. 2000; Decreto-Lei n° 92/2019. 2019). Coastal species and NIS have been usually monitored through observational and morphotaxonomic approaches. With the progress of sequencing technology, DNA-based methods are becoming part of monitoring efforts and gradually implemented in formal national monitoring programmes (Weigand et al. 2019). Indeed, previous studies indicate that metabarcoding delivers results comparable to morphological surveys, with similar implications for management actions, at a considerably lower cost (Borrell et al. 2017; Ji et al. 2013; Rey et al. 2020). The use of DNA metabarcoding has many advantages, including the possibility to screen numerous species simultaneously. However, for a more robust identification of NIS, species detected by molecular methods should be confirmed through morphological identification to prevent false positives (Fonseca et al. 2023) or using more targeted molecular approaches (e.g. qPCR, ddPCR) (Ammon et al. 2018; Wood et al. 2019). To the best of our knowledge, official NIS lists, such as those released by the International Council for the Exploration of the Sea (ICES) and the Convention for the Protection of the Marine Environment of the North-East Atlantic (OSPAR) commission, for example, are opting to include identifications obtained through environmental DNA only if they were also supported by morphological identifications. This is due to the limitations of DNA metabarcoding identifications that include the incompleteness of reference libraries; operational limitations (false positives and negatives, primer bias and contamination) and the lack of standardization (Ruppert et al. 2019; Duarte et al. 2021), as well as the inability to generate absolute abundance data and population status. Other complications may arise due to the inaccuracy or ambiguity of the reference sequences available in publicly-accessible databases (Fontes et al. 2021; Radulovici et al. 2021; Lavrador et al. 2023), which made us to exclude some NIS records from this study that had uncertain species identifications. For example, *Watersipora subtorquata*, a well-known and highly invasive NIS, was only detected (using the COI marker) in BINs assigned to more than one species, that could not be solved by our curation method. This was also the case for *Botrylloides violaceus* and *Lovenella assimilis* as previously mentioned, that were only detected using 18S due to the inaccurate matching records in the COI reference data. However, in the case of *W. subtorquata*, studies have shown that several clades form a complex of cryptic species that make molecular identification more difficult or even unattainable until the status of the MOTUs of the complex is clarified (MacKie et al. 2012; Duncan et al. 2022). These still existing problems of lack of completion and uncertain accuracy of certain records in typically used databases were also noticeable in our study, as when curating NIS molecular detections with both markers, only 27% (18S) to 42% (COI) of these records were considered reliable. These challenges highlight the need for further research to address taxonomic uncertainty issues and enhance the reliability of DNA metabarcoding as a tool for NIS identification.

Many efforts have been made recently to publish updated lists of NIS in European waters and to further monitor its distribution as well as to study its introduction history (Gittenberger et al. 2023; Jensen et al. 2023; Png-Gonzalez et al. 2023). In a recent NIS inventory in Spanish marine waters, approximately 65% of NIS were invertebrates (Png-Gonzalez et al. 2023). In Danish waters, 40% of NIS were invertebrates comprising both benthic invertebrates and zooplankton (Jensen et al. 2023). In a study of NIS in the coastal waters of the Netherlands, the majority of invertebrate NIS belonged to Crustacea, Mollusca, and Annelida. With a mention that bryozoans have become a very relevant group of NIS, which may be connected to hull fouling as one of the most important vector in NIS introduction in recent years (Gittenberger et al. 2023). The same study also concluded that the number of introductions by biofouling appears to have increased in the last decade, while introductions by aquaculture and fisheries seem to have decreased (Gittenberger et al. 2023). In addition, it is very common to observe detached gear and other floating litter, namely plastic, in ports and recreational marinas. Marine plastic litter can also provide a surface for organisms’ colonization and establishment, facilitating its spread to new locations (Barnes 2002; Ibabe et al. 2020). Some studies have shown the ability of biota to experience long trans-oceanic transport and to survive over years attached to floating litter (Therriault et al. 2018). This increasing interest in studying marine NIS, particularly in recreational marinas and ports, will help to understand the best methodologies to monitor these species, to potentially reach a global standardized protocol for detection of marine invertebrates NIS in these environments.

Although the use of DNA-based tools for monitoring marine invertebrate NIS in marinas and ports still needs further refinement, its exceptional potential is already apparent. The employment of these techniques constitutes an important complement or early warning system that will enable to manage in a timelier manner the detection of an invasive species at an early stage of introduction. For example, in the Port of Vlissingen, the genetic material of a NIS was detected three years before the specimens were physically found (Gittenberger et al. 2023). Early detection of NIS is also important to determine pathways and vectors of introductions, as well as to resolve cryptic species (Gittenberger et al. 2023). This study demonstrated that to monitor NIS in recreational marinas, a combined approach is required (in order to recover a broader spectrum of taxa and to provide a comprehensive picture of the surveyed species and communities). To this end, it is indispensable to employ not only several genetic markers, but also to consider different sampling procedures, a variety of target substrates and seasonal variation in the sampling design.

## Materials and Methods

### Study sites

Sampling was conducted in two recreational marinas located in the Northwest of Portugal: the recreational marina of Viana do Castelo (Latitude: 41°40.5’N, Longitude: 8°50.3’W) and the Porto Atlântico marina (Latitude: 41°11.0’N, Longitude: 08°42.3’W) (Fig. 6). The recreational marina of Viana do Castelo is located in the north bank of the Lima River estuary, in the city of Viana do Castelo, and has a capacity for docking 300 vessels, a maximum length of 20 m and a depth of ca. 3 m, usually maintained by dredging (Porto de Viana do Castelo 2017). Several hundreds of boats dock at the recreational marina annually, the majority from France, the United Kingdom, the Netherlands, and Germany (Porto de Viana do Castelo, 2017), but overall, it presents a small ship traffic, since it is located further upstream, where commercial and fishing ports from Viana do Castelo are located.

**Fig. 6.**
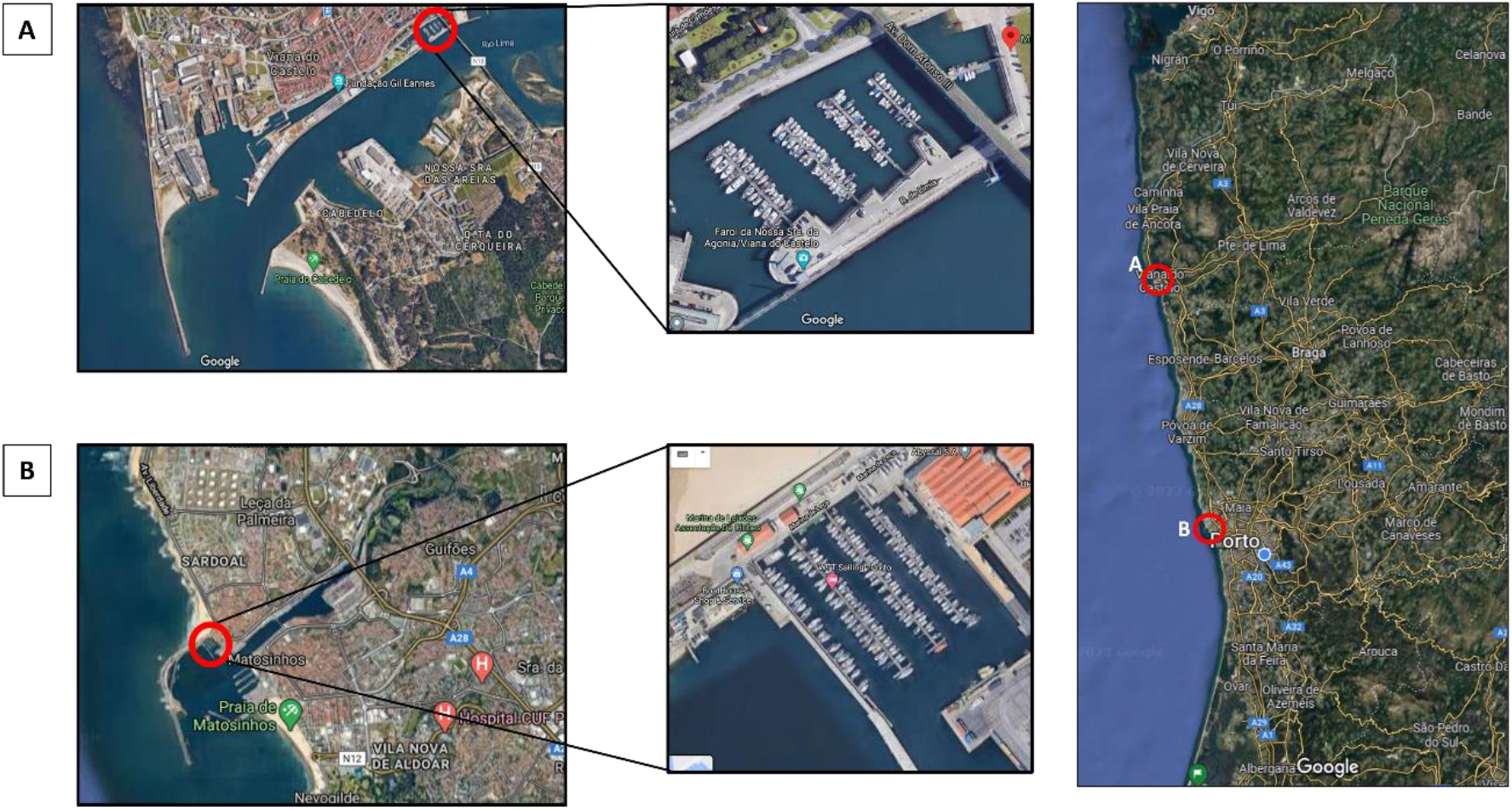
Images retrieved from Google Maps showing on the left: the location of the two sampled recreational marinas and their proximity to the ocean; on the right: the northern region of Portugal containing the location of the two marinas, which are separated by a distance of approx. 56.8 km. **A** Viana de Castelo. **B** Porto Atlântico

The Porto Atlântico is an artificial marina with a capacity for docking 240 vessels, a maximum length of 35 m and a depth that can vary between 2 to 4 m. It is located 2.5 miles from the mouth of the Douro River, in Matosinhos and at the entrance of the Leixões harbour. The Porto Atlântico harbour is the second most relevant at national level in terms of merchandise movement with 15 188 000 tons being handled in this harbour in 2021, of which ca. 79% was from international commerce (APDL, 2021). In 2021, Viana do Castelo harbour received a total of 250 ships, while Porto Atlântico harboured a total of 2,410 ships (APDL, 2021).

Sampling was conducted in 3 different seasons – spring (T1), autumn (T2) and winter (T3), in 3 different points of each marina, to obtain a larger spatial representativity. On each sampling date, physical and chemical parameters, namely conductivity, salinity and pH were measured using a multiparameter probe (Multiline F/set 3 no. 400327, WTW, Weilheim, Germany) (Table S5). Water temperature data were obtained from the Portuguese Institute for the Sea and Atmosphere (IPMA, I.P.) (https://www.ipma.pt/pt/maritima/costeira/), as well as data from the vessels on dock at each harbour closest to the recreational marinas, from the Administration of Douro, Leixões and Viana do Castelo Harbours (APDL) (https://viana.apdl.pt/ and https://leixoes.apdl.pt/ for Viana do Castelo and Porto Atlântico respectively), during the entire experiment time (March 2020 to March 2021).

### Sample types

On each sampling date, 5 strategies were employed: i) organisms fouling into hard substrates of each artificial marina; ii) organisms colonizing sponges (artificial substrates) and iii) organisms colonizing acrylic plates (artificial substrates) that were deployed on each marina, during fixed periods of time (4-5 months); iv) water collection for environmental DNA (eDNA) analysis and v) zooplankton collection from the water column.

### Hard substrates

On each sampling point, within each marina, the organisms fouling the marina’s hard substrates, such as pontoons, cables, ropes, and buoys, were scraped from a total area of 22 x 22 cm, into zip-lock bags, and stored at 4 °C with water from each site, until processing. In the lab, the samples were poured through a 500 µm mesh sieve (previously washed with 10% bleach), with the water of each respective bag, and the organisms retained in the sieve were stored in containers with absolute ethanol, at -20 °C, until DNA extraction.

### Artificial substrates

Two types of artificial substrates were deployed on each sampling point: i) tri-dimensional nylon bath sponges, with 1 mm-mesh size, which can retain different organisms, as well as act as a substrate to sessile organisms and ii) white acrylic plates (similar to the material of the submerged part of the pontoons that was scraped) of 22 x 22 cm. These substrates (Fig. 7) were deployed on the 2nd of March 2020 on each point from each marina, close to the pontoons and with a depth of 1 m and collected and processed in the sampling dates above-mentioned. Due to the restrictions imposed by the COVID-19 outbreak, the artificial substrates sampled in the winter were deployed in the water column for longer (140 days total) than those sampled in spring (105 to 107 days) and autumn (119 days). On each sampling date, each artificial substrate was placed in separate zip-lock bags and stored at 4 °C, with water from each site, and new artificial substrates were deployed until the next sampling season. In the lab, the organisms fouling the tri-dimensional sponges as well as the water of each respective bag were thoroughly washed through a 500 µm sieve (previously washed with 10% bleach and MilliQ water). The plates were scraped and also washed through a 500 µm sieve. The organisms retained in the sieve were stored in containers with absolute ethanol, at -20 °C, until DNA extraction.

**Fig. 7.**
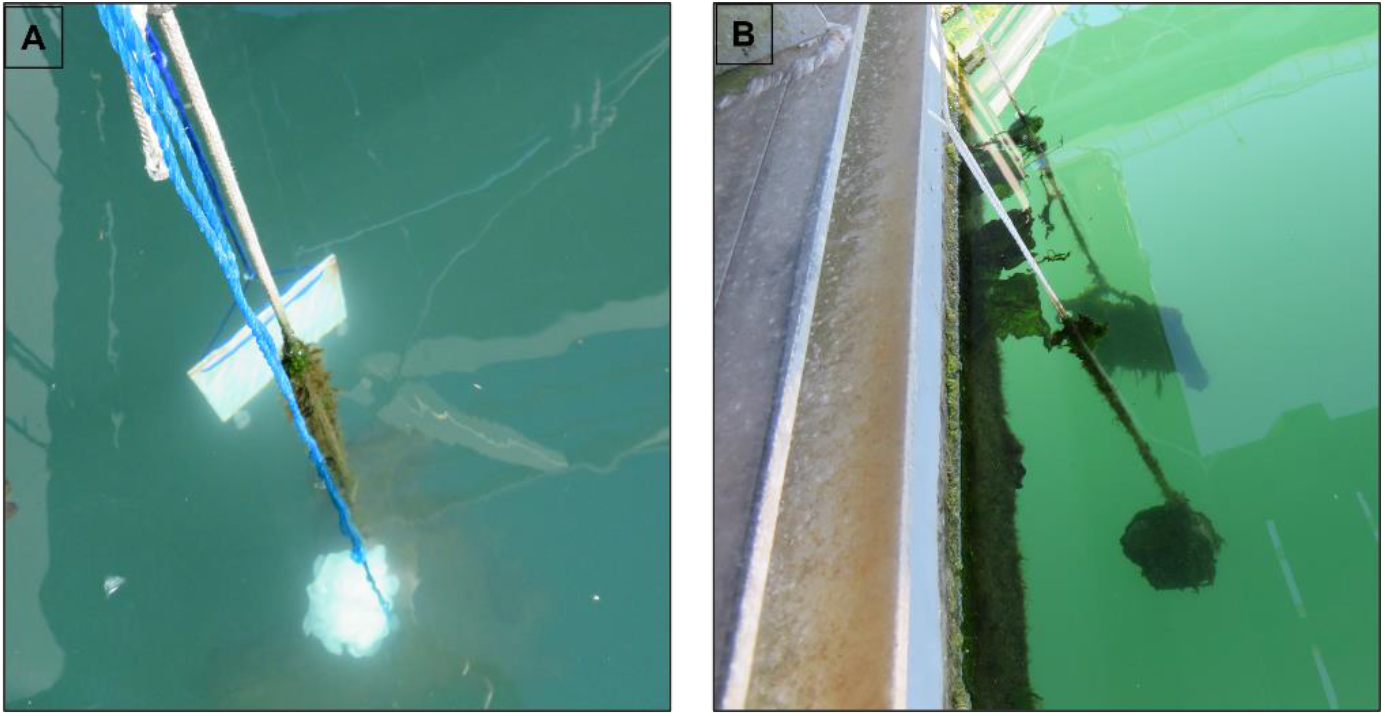
Pictures showing the employment of the artificial substrates. **A** Artificial substrates at the beginning of a sampling season. **B** Artificial substrates after colonization.

### eDNA

One litre of water at ca. 1 m depth was collected using a cable, close to the pontoons, in polypropylene flasks, previously washed with 10% bleach and rinsed with MilliQ water. The cable was also previously cleaned with 10% bleach, before immersion at each sampling point. On each sampling point, each flask was washed 3 times with the water of the respective site. After collection, the water was stored at 4 °C for a maximum period of 24 h, filtered through a 0.45 µm pore size filter (S-Pak Filters, Millipore) and filters were then stored at -20 °C, until DNA extraction. Negative controls, consisting in one 1L flask, containing MilliQ water, was used throughout all the workflow (sampling, storage, sample processing in the lab, DNA extraction and PCR amplification) and processed as the eDNA samples.

### Zooplankton

Samples were obtained using a plankton net with 55 µm mesh size, with a mouth of 40 cm of diameter and 100 cm length. On each sampling point, 3 separate oblique tows were performed for 45 seconds each and the end-cup content was poured into 1L polypropylene flasks, also previously washed with 10% bleach and MilliQ water. After collection, the concentrated samples were stored at 4 °C for a maximum period of 24 h and filtered through a 0.45 µm pore size filter (S-Pak Filters, Millipore) and filters were stored at -20 °C, until DNA extraction.

### DNA extraction

The DNA from the hard and artificial substrates was extracted using a silica-based method, adapted from Ivanova et al. (2006), and as described by Steinke et al. (2022). Briefly, up to 30 g of ethanol-preserved organisms were placed in autoclaved flasks, previously washed with 10% bleach and Milli-Q water, to which an adequate volume of a Lysis Buffer (100 mM NaCl, 50 mM Tris-HCl pH 8.0, 10 mM EDTA pH 8.0 and 0.5% SDS) (depending on the sample wet weight), was added. Samples were then digested overnight in an orbital incubator (Infors MT Multitron Pro) at 40 rpm and 56 °C. The lysates were then centrifuged, and supernatants mixed with a Binding Mix (6M GuSCN, 20mM EDTA pH 8.0, 10mM Tris-HCl pH 6.4 and 4% Triton X-100) and purified through silica columns following 3 washing steps, with two ethanol-based solutions (Protein Wash Buffer and Wash Buffer). DNA was finally eluted from the columns using autoclaved deionized water.

Genomic DNA from eDNA and zooplankton samples was extracted from half of the filters using the DNeasy PowerSoil Kit (Qiagen), following the manufacturer’s instructions.

Negative controls were processed along the DNA extraction procedure to check for contaminations of the solutions and labware materials used. These negative controls were used as templates in subsequent PCR amplification reactions.

### Amplicon libraries and high-throughput sequencing (HTS)

Amplicon libraries and high-throughput sequencing (HTS) were carried out at Genoinseq (Biocant, Portugal). To amplify the internal region of 313 bp of the mitochondrial cytochrome c oxidase I (COI) gene, the primer pair mlCOIintF (5’-GGWACWGGWTGAACWGTWTAYCCYCC - 3’) (Leray et al. 2013) and LoboR1 (5’-TAAACYTCWGGRTGWCCRAARAAYCA - 3’) (Lobo et al. 2013) was used, while the primer pair TAReuk454FWD1 (5’-CCAGCASCYGCGGTAATTCC - 3’) and TAReukREV3 (5’-ACTTTCGTTCTTGATYRA - 3’) (Stoeck et al. 2010; Lejzerowicz et al. 2015) was used to amplify the ∼400 bp of the V4 hypervariable region of the 18S rRNA gene (18S). The two primer pairs were selected based on previous studies on marine invertebrates of the region (Duarte et al., 2023; Fais et al., 2020; Leite et al., 2021). For each sample, PCR reactions were performed using KAPA HiFi HotStart PCR kit according to manufacturer instructions. For the COI gene amplification, the PCR conditions involved a 3 min denaturation at 95°C, followed by 35 cycles of 98°C for 20 s, 60°C for 30 s and 72°C for 30 s and a final extension at 72°C for 5 min. For the 18S gene, the PCR conditions involved a 3 min denaturation at 95°C, followed by 10 cycles of 98°C for 20 s, 57°C for 30 s and 72°C for 30 s and 25 cycles of 98°C for 20 s, 47°C for 30 s and 72°C for 30s, and a final extension at 72°C for 5 min.

Second limited-cycle PCR reactions added indexes and sequencing adapters to both ends of the amplified target regions according to manufacturer recommendations (Illumina 2013). For purification of the PCR products, the SequalPrep 96-well plate kit was used (ThermoFisher Scientific, Waltham, USA) (Comeau et al. 2017). The products were then pooled, and pair-end sequenced in the Illumina MiSeq® sequencer with the V3 chemistry, according to manufacturer instructions (Illumina, San Diego, CA, USA) at Genoinseq (Biocant, Portugal).

### Bioinformatics pipelines

The raw reads (in fastq format) obtained from the Illumina MiSeq® System were subjected to quality filtering using PRINSEQ version 0.20.4 (Schmieder and Edwards 2011). This involved the elimination of sequencing adapters and reads with less than 100 bp for COI and less than 150 bp for 18S. Additionally, bases with an average quality below Q25 within a 5-bases window were trimmed. The mothur software (version 1.39.5) was used to merge the filtered forward and reverse reads, by overlapping paired-end reads (make.contigs function, default alignment). Primer sequences were also removed during this step using the trim.seqs function (default settings) (Schloss et al. 2009; Kozich et al. 2013).

Two database pipelines were used to process the reads: mBrave – Multiplex Barcode Research and Visualization Environment (www.mbrave.net; Ratnasingham, 2019), which is connected to BOLD (Ratnasingham and Hebert 2007), was used for the COI reads; while the 18S reads were analyzed in the SILVAngs database (https://ngs.arb-silva.de/silvangs/; Quast et al., 2013).

In mBrave, the COI reads were uploaded using the sample batch function, and only length trimming was applied, with a maximum length set at 313 bp. Subsequently, low-quality reads were discarded based on two criteria: average quality value (QV) less than 10 and sequences shorter than 150 bp. Reads satisfying these criteria underwent de-replication, followed by the removal of chimeras. The remaining reads were clustered into Operational Taxonomic Units (OTUs) using a distance threshold of 3%. Taxonomic assignment at the species level using a 97% similarity threshold was performed for the resulting OTUs by comparing them against the BOLD database, as well as against several public datasets specific for marine invertebrates of the Northeast Atlantic previously published: DS-GAIMARIN (Leite et al., 2020), DS-BIBLIO (Lobo et al., 2017), DS-PTGB (Lobo et al. 2016), DS-3150 (Lobo et al. 2017b), DS-PMACA (Vieira et al. 2022) and DS-NISEUREF (Lavrador et al. 2023). The Barcode Index Number (BIN) system englobe a cluster of COI nucleotide sequences to produce operational taxonomic units that usually represent a species molecularly, using well established algorithms in BOLD (Ratnasingham and Hebert 2013). In mBrave, HTS data is matched to these BINs whenever possible, to obtain a taxonomic identification of the sequences to species level, but for some cases a further audition and curation is needed. These matches were audited with the following methodology: 1) matches with BINs attributed to more than one species were identified and flagged; 2) those matches were audited using the protocol detailed in Lavrador et al., 2023. Briefly, matches with BINs attributed to 3 or more morphospecies were automatically discarded. On the other hand, BINs attributed to 2 morphospecies were analysed for cases of synonyms or misidentification that could be solved. If these cases of discordances had resolution, taxonomic matches would remain for further analysis, otherwise the taxonomic assignment to species level was automatically discarded.

In SILVAngs, the SILVA Incremental Aligner (SINA v1.2.10 for ARB SVN (revision 21008)) (Pruesse et al. 2012) was used to align each read against the SILVA SSU rRNA SEED database and perform quality control (Quast et al. 2013). Criteria for exclusion were reads with less than 150 aligned nucleotides and with more than 1% ambiguities or 2% homopolymers. Reads identified as putative contaminations or artifacts, based on low alignment quality (80 alignment identity, 40 alignment score reported by SINA), were also excluded from downstream processing. VSEARCH (version 2.14.2; https://github.com/torognes/vsearch) (Rognes et al. 2016) was used for dereplication and cluster of the unique reads in OTUs per sample, applying identity criteria of 1.00 and 0.99, respectively. After these initial steps of quality control, identical reads were identified (dereplication), the unique reads were clustered (OTUs) on a per sample basis, and then the reference read of each OTU was taxonomically assigned. Taxonomic assignment was performed using BLASTn (2.2.30+; http://blast.ncbi.nlm.nih.gov/Blast.cgi) (Camacho et al. 2009) with standard settings, against the non-redundant version of the SILVA SSU Ref dataset (release 138.1; http://www.arb-silva.de). The taxonomic classification of each OTU reference read was applied to all reads assigned to that respective OTU. Reads with weak or no classifications, with a “(% sequence identity + % alignment coverage)/2” value below 70, were assigned to the category “No Taxonomic Match.” For subsequent analysis, only OTUs taxonomically identified with a similarity threshold of 99% were retained. NIS detected exclusively with this marker were further analysed by performing a separate BLASTn with standard settings, against standard databases of NCBI, to verify the reliability of the taxonomic assignments.

For both markers, only reads assigned species level and that belonged to metazoan invertebrate groups were used for further analysis, since in the case of non-indigenous species, information at species level is mandatory for better tailoring an effective management strategy. Species taxonomic classification and environment were verified in the World Register of Marine Species (WoRMS) database (www.marinespecies.org, accessed on 27th April 2023) (World Register of Marine Species, 2023). Only marine or oligohaline invertebrate species were retained.

For the identification of NIS, final species lists were matched to Portuguese invertebrate updated NIS lists (gently shared with us by Paula Chainho, which is the Portuguese contact point for the Working Group on Introductions and Transfers of Marine Organisms (WGITMO) of the ICES) as well as to an updated European NIS list. The European NIS list was obtained from three public databases: the European Alien Species Information Network (EASIN) (https://easin.jrc.ec.europa.eu/easin, accessed on 22nd March 2023) (Katsanevakis et al. 2012), the Information System on Aquatic Non-indigenous and Cryptogenic Species (AquaNIS) (http://www.corpi.ku.lt/databases/index.php/aquanis/, accessed on 17th March 2023) (Olenin et al. 2014) and the World Register of Introduced Marine Species (WRiMS) (https://www.marinespecies.org/introduced, accessed on 22nd March 2023) (Rius et al. 2022) (as described in Lavrador et al., 2023).

### Statistical analyses

Venn diagrams were generated to determine the overlap between species and NIS detected with COI and 18S, as well as between species and NIS detected on each sample type, detected on each season and detected at each location (https://bioinformatics.psb.ugent.be/webtools/Venn/) were used to display the distribution of species and NIS among phyla, for each sampled marina, using GraphPad Prism v8 (GraphPad Software, Inc.).

Principal coordinate analysis (PCoA), using the Jaccard index, were performed to visualize the similarity on community structure among the different sample types and seasons, for each marina. A permutational variance analysis (PERMANOVA), using 999 permutations, was then used to test the effect of the factor’s “season” and “sample type” on the community structure of the marine invertebrates recovered. PCoA and the PERMANOVA analysis were performed on Primer v6.1.16 (Clarke and Gorley 2006).

## Acknowledgements

We would like to thank Paula Chainho, Romeu Ribeiro and João Gil for their help with identifying species as non-indigenous for Portugal. This work was funded by the project “NIS-DNA: Early detection and monitoring of non-indigenous species (NIS) in coastal ecosystems based on high-throughput sequencing tools” funded by the Portuguese foundation of Science and Technology (FCT, I.P. under the reference PTDC/BIA-BMA/29754/2017) and by the “Contrato-Programa” (https://doi.org/10.54499/UIDB/04050/2020), funded by national funds through the FCT I.P. https://doi.org/10.54499/UIDB/04050/2020. Financial support granted by the FCT I.P. to S.D. (https://doi.org/10.54499/CEECIND/00667/2017/CP1458/CT0001) is also acknowledged. A.S.L. (UI/BD/150871/2021) is supported by the Collaboration Protocol for Financing the Multiannual Research Grants Plan for Doctoral Students with financial support from FCT I.P. and the European Social Fund under the Northern Regional Operational Program—Norte2020.

## Author Contributions

SD and FOC conceptualized, administered and acquired funding for the study; ASL, SD, PEV, FGA and JM performed the sampling and laboratory procedures; ASL, SD and FOC analysed the data and discussed the results; ASL and SD wrote the paper; ASL, FGA, JM, PEV, FOC and SD revised and reviewed the paper. All authors have read and agreed to the published version of the manuscript.

## Declarations

### Conflict of interest

The authors declare that they do not have any conflict of interest.

## Notes

### Competing Interest Statement

The authors have declared no competing interest.

