## Supplemental Figures for "DNA metabarcoding-based detection of non-indigenous invertebrates in recreational marinas: influence of sample type and seasonal variation"

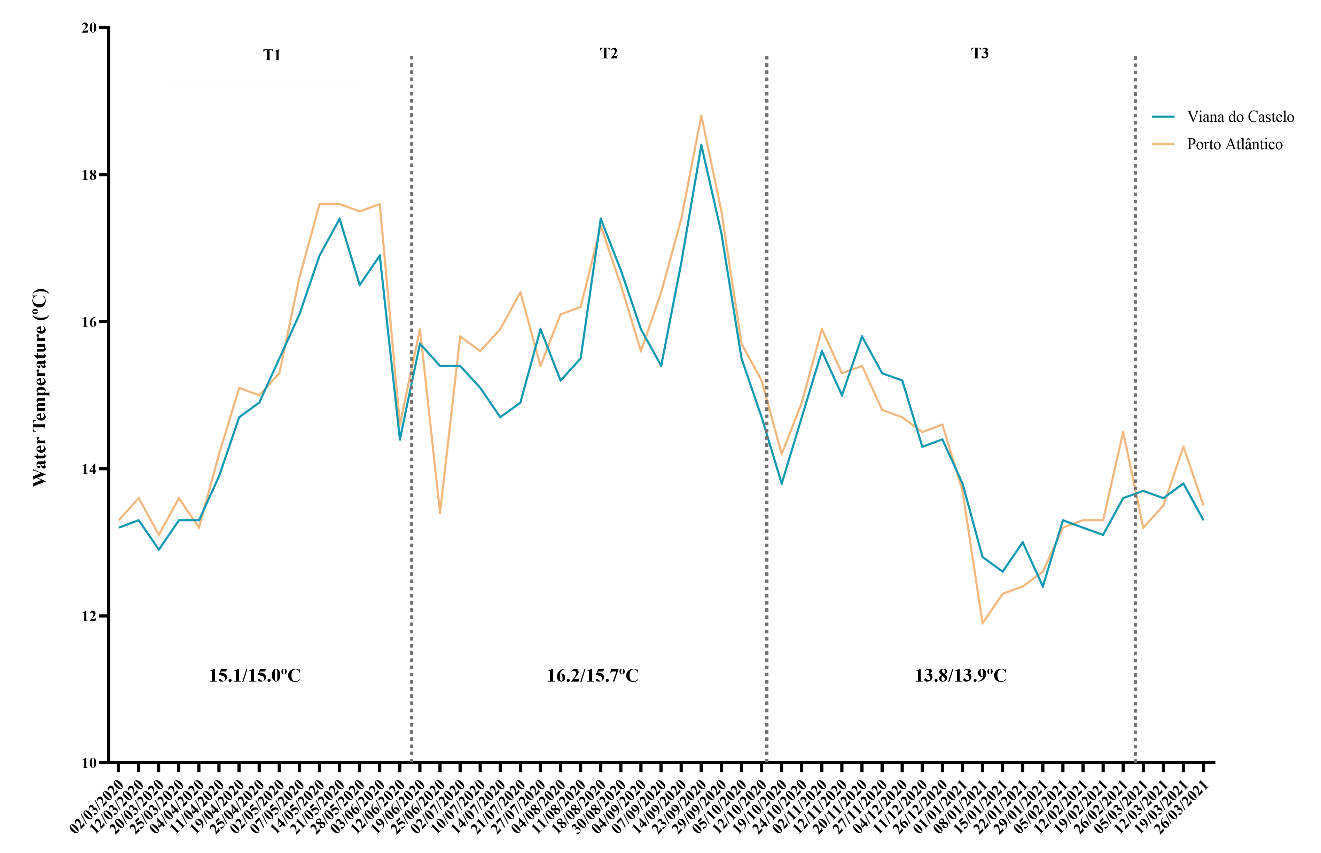

**Figure S1**. Variation of water temperature recorded throughout the experimental time in the beaches closest to each of the recreational marinas. Dotted lines indicate sampling dates, and separate the data in the graph in seasons. Numbers on the bottom of the graph represent the average temperature for each season and location (Porto Atlântico/Viana do Castelo).

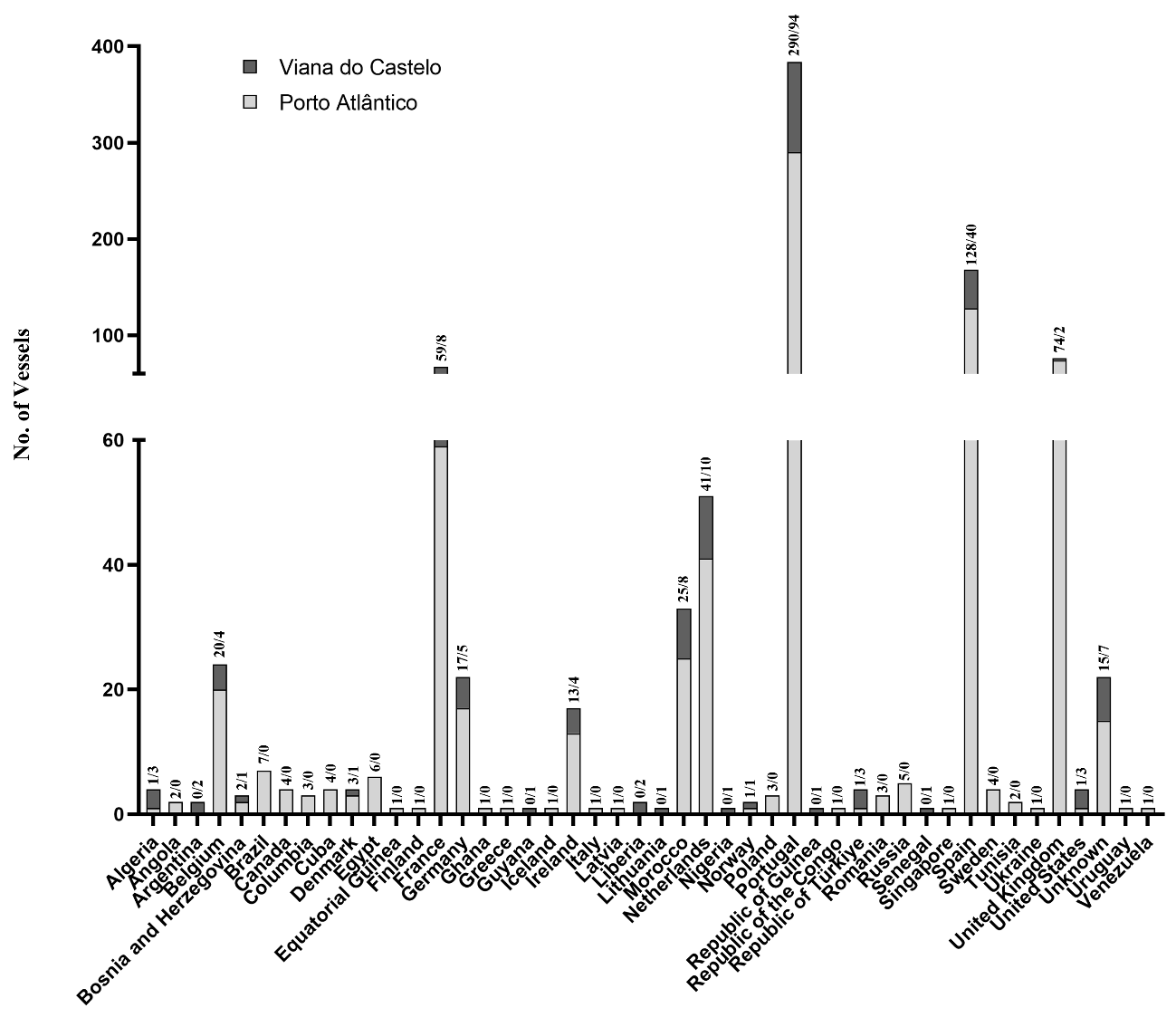

**Figure S2**. Number of Vessels per country of origin and per harbour closest to each of the sampled recreational marinas. On top of each column bar are represented the total number of vessels per harbour (Porto Atlântico/Viana do Castelo).
